# Mutant ASXL1 Drives Transcriptional Activation and Repression in Human Hematopoiesis

**DOI:** 10.64898/2026.01.13.699315

**Authors:** Madison L. Hall, Aliya Quintal, Samantha Worme, Thai Nguyen, Zachary Schonrock, Mitsuhiro Tsuchiya, Sonia N. Acharya, Hanqian L. Carlson, Itallia V. Pacentine, Shawn Shrestha, Jommel Macaraeg, Randall Armstrong, James McGann, Sara Evans-Dutson, Theresa A. Lusardi, Brendan L. O’Connell, Andrew C. Adey, Galip Gürkan Yardimci, Julia E. Maxson, Theodore P. Braun

## Abstract

Mutations in the epigenetic regulator *ASXL1* are common in myeloid malignancies and portend a near-universally poor prognosis. While multiple mechanisms for mutant ASXL1-dependent oncogenesis have been proposed, none have been functionally validated in the context of the human hematopoietic stem cell, where these mutations almost certainly arise. Here, we extensively characterized a CRISPR-engineered human hematopoietic stem and progenitor cell model of ASXL1 mutations. In this context, mutant ASXL1 expression decreases differentiation, increases clonogenicity in serial replating experiments, and improves engraftment in immunodeficient mice. We also show that endogenous truncating mutations in ASXL1 drive protein stabilization and confirm that mutant ASXL1 is resistant to proteasomal degradation. At the transcriptional level, these phenotypes are driven by significant repression of the stress-response genes and by increased expression of bromodomain and extra-terminal family protein targets. Using protein-interaction screens, genomic and functional approaches, we link the positive transcriptional changes in *ASXL1*-mutant cells to BRD4-dependent RNA polymerase II pause release and identify a mechanism for transcriptional repression via a previously uncharacterized interaction with the transcription factor MECOM. Finally, we demonstrate that *ASXL1*-mutant AML exhibits increased MECOM activity consistent with our gene-editing models. Collectively, these studies highlight a highly reproducible model of mutant *ASXL1* in the appropriate cell context. Further, they are the first to functionally describe the mutant *ASXL1* interactome in the context of the human HSC, identifying new dependencies with therapeutic potential.

## Introduction

Mutations in the epigenetic regulator ASXL1 are early genetic events in myeloid malignancies and are associated with high rates of progression and treatment resistance across the disease spectrum^1–4^. These mutations are the third most common in the pre-malignant condition age-related clonal hematopoiesis, where they portend a 50% risk of progression to overt malignancy over 10 years^1^. In patients with myelodysplastic syndrome (MDS), ASXL1 mutations are associated with a 50% reduction in overall survival (19 mo vs 39 mo) and an increased rate of transformation to acute myeloid leukemia (AML, HR 2.2)^5,6^. Once AML develops, mutant ASXL1 continues to drive decreased overall survival (15.9 mo vs 23 mo)^3^. We lack effective clinical therapies that target mutant ASXL1 or its downstream dependencies, representing a significant unmet need.

The function of wild-type and mutant ASXL1 remains incompletely understood. Early studies in Drosophila found both activating and repressive functions for ASXL1, with gene disruption producing both anterior and posterior homeotic defects^7,8^. While the phenotype of ASXL1 disruption alone is relatively mild, it enhances the phenotype of trithorax (activating histone methyltransferase) and polycomb (PRC, deposits repressive histone modification) group mutations, leading to designation as an enhancer of trithorax and polycomb (ETP) family protein. In the hematopoietic system, deletion of ASXL1 produces only minimal changes in the landscape of covalent histone modifications, instead driving changes in RNA polymerase II localization^9^. However, the relevance of this finding to myeloid malignancies remains unclear.

Disease-associated mutations in *ASXL1* result in the production of a truncated protein with altered activity^10–13^. Multiple mechanisms have been proposed to explain the impact of mutant ASXL1. The best studied of these is hyperactivation of the BAP1 deubiquitinase, leading to the removal of PRC1-deposited repressive H2aK119Ub and gene activation^10,11,13^. However, BAP1 also deubiquitinates mutant ASXL1 directly, thereby stabilizing the protein and increasing its chromatin binding^13^. Meanwhile, mutant ASXL1 also interacts with the transcriptional activator BRD4, driving gene activation^12,14^. It is unclear how these mechanisms interact to drive phenotypic changes in ASXL1 mutant myeloid cells.

Here, we demonstrate that mutant ASXL1 is highly localized to active transcriptional complexes and alters RNAPII recruitment. To functionally validate these findings, we developed a high-efficiency CRISPR-engineered model of *ASXL1* mutations in human CD34+ cells. These cells resist differentiation, exhibit increased clonogenicity, and show enhanced engraftment in immunocompromised mice. Using proximity-dependent biotin ligation and an arrayed CRISPR screen, we found that mutant ASXL1 interacts with and depends upon multiple proteins in active transcriptional complexes, including members of the FACT and ISWI complexes as well as BRD4. We also identified a previously unappreciated interaction with the oncogenic transcription factor MECOM, which is typically a transcriptional repressor^15,16^. Mutant ASXL1 co-localizes with MECOM at FOS and FOSB, genes that drive HSC quiescence and promote self-renewal. MECOM knockout reverses the ability of mutant ASXL1 to increase clonogenicity and resistance to differentiation in vitro and increase engraftment in vivo. We also show that MECOM activity is increased in *ASXL1*-mutant chronic myelomonocytic leukemia and AML, thereby establishing the validity of this novel mechanism of mutant ASXL1-driven oncogenesis.

## Results

### Mutant ASXL1 Co-Localizes with Active Transcriptional Complexes

**—**Mutant ASXL1 is highly co-bound with RPB1, the large subunit of RNAPII (**Figure 1A, B, C**)^10^. In contrast, mutant ASXL1 binding sites are almost entirely separate from H2aK119Ub domains, though there are rare instances where RPB1, ASXL1, and H2K119Ub co-localize (**Figure 1B**). This argues that the direct transcriptional effects of mutant ASXL1 depend on BAP1-driven ASXL1 protein stabilization rather than direct changes in H2aK119Ub. Both mutant and wild-type ASXL1 bind primarily at promoter regions, suggesting that ASXL1 may play a direct role in facilitating the increased RNAPII pause-release in our prior work. (**Figure 1D**)^9^. However, transcriptional regulation is highly context-dependent, and cell lines may not accurately model ASXL1 mutations in hematopoietic stem cells, where they drive clonal expansion early in disease evolution.

**Figure 1.**
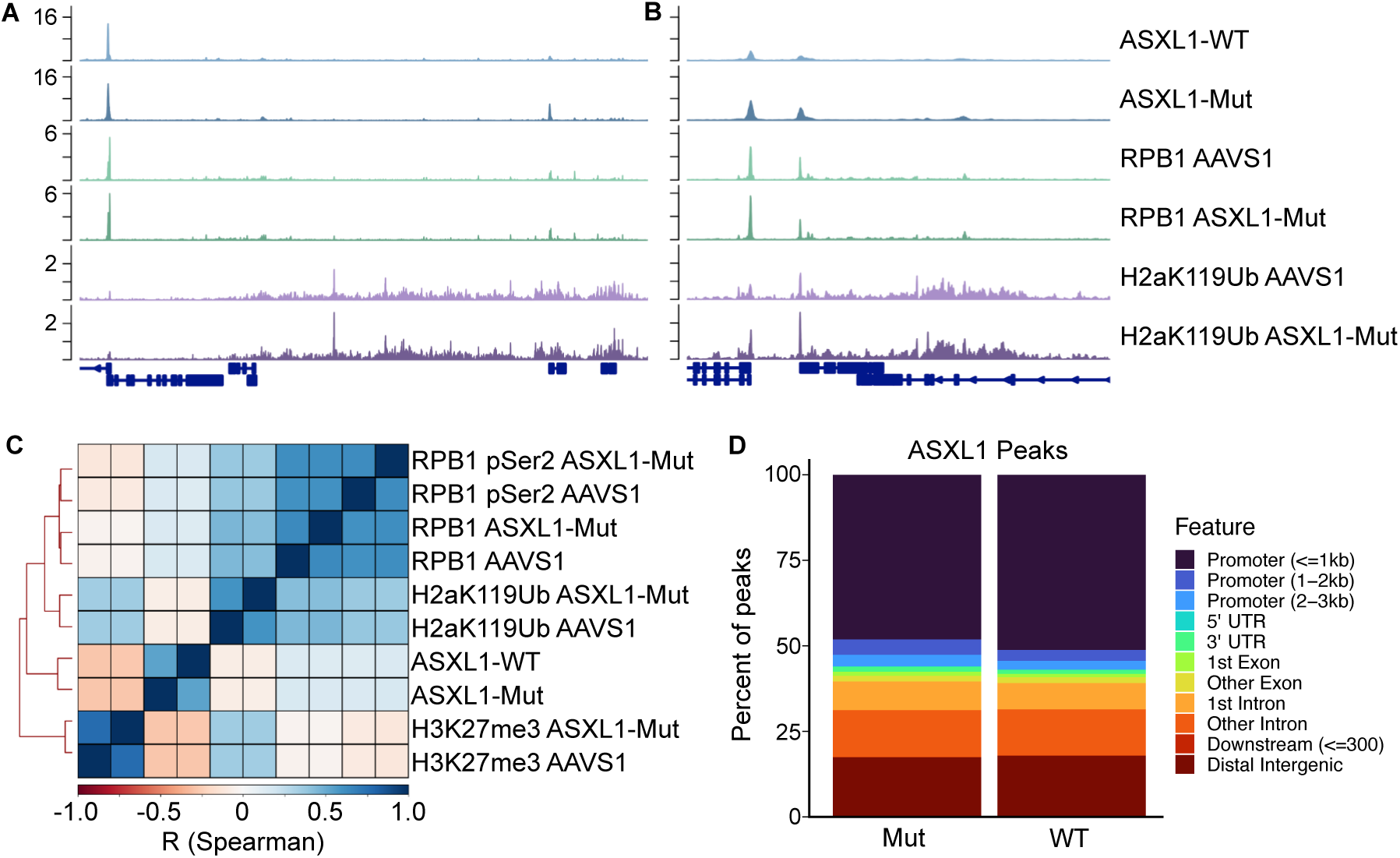
Mutant and Wild-Type ASXL1 Co-Localize with Active Chromatin. **A, B**. ChIP-seq for ASXL1 from *ASXL1*-mutant and wild-type cells overlayed with CUT&Tag for RPB1 (large subunit of RNAPII) and H2aK119Ub shows colocalization of ASXL1 with RNAPII and occasional co-localization with H2aK119Ub. **C.** Genome-wide Spearman correlation of data in **A** and **B.**

### Modeling ASXL1 Mutations in Human Hematopoietic Cells

**—**To study mutant ASXL1 in hematopoietic stem and progenitor cells, we developed a CRISPR gene-editing strategy using triplet sgRNAs, resulting in truncating mutations in *ASXL1* at nearly 100% efficiency (**Figure 2A and Supplementary Figure 1A**). When cultured under pro-differentiation conditions, *ASXL1*-mutant HSPCs resisted differentiation, thereby retaining cells in the CD34+ compartment (**Figure 2B**). Furthermore, *ASXL1* mutant cells exhibited increased replating capacity in a colony-forming assay (**Figures 2C and 2D**), with increased colony output intermittently seen in the initial round of plating (i.e., **Figure 2G**). In contrast, targeting the second exon of *ASXL1* with sgRNAs (resulting in a functional knockout) increased differentiation and decreased replating capacity. Upon transplantation into immunodeficient mice, *ASXL1*-mutant cells exhibit increased engraftment and myeloid-biased output (**Figure 2E**).

**Figure 2.**
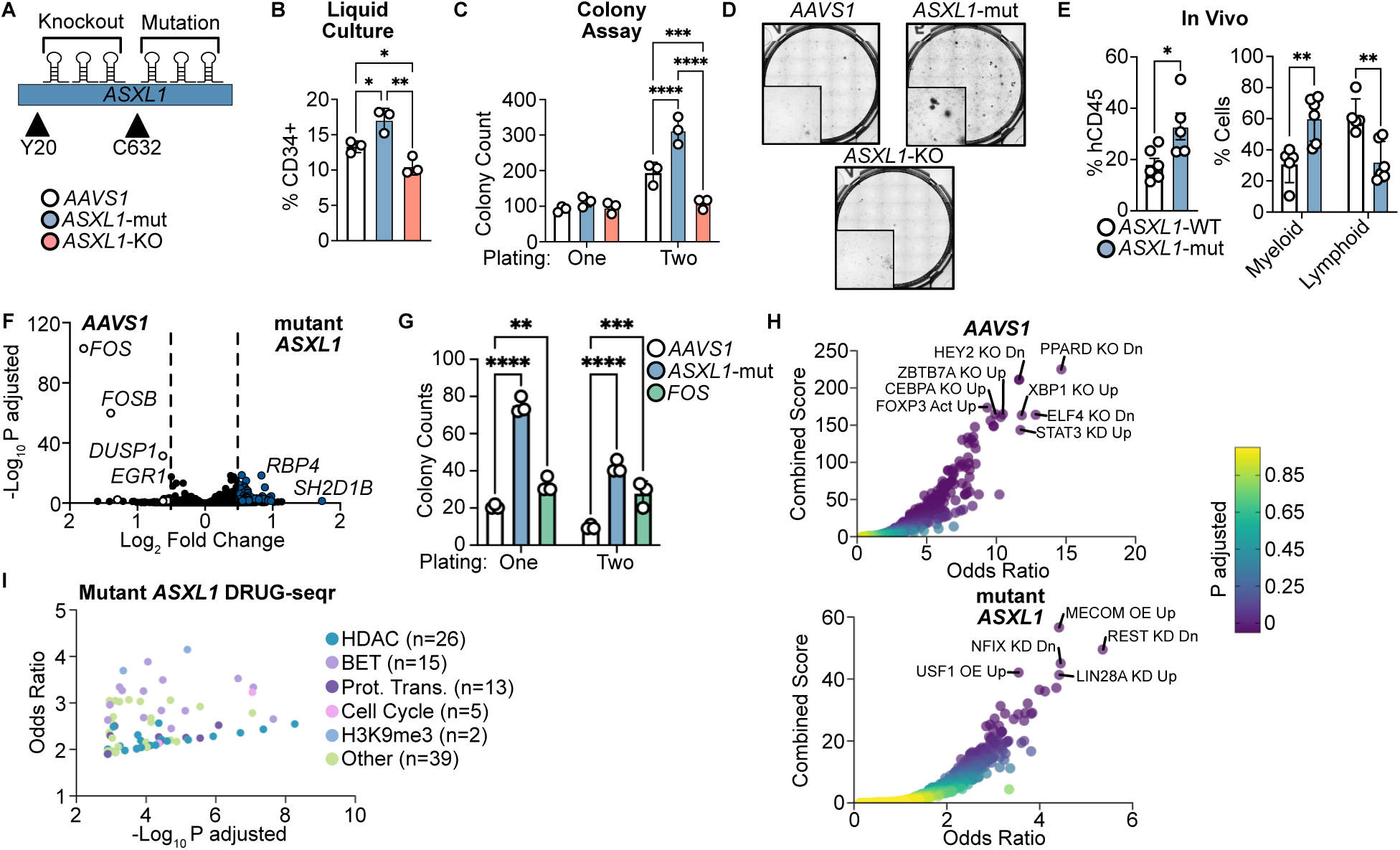
Modeling *ASXL1* Mutations in CD34+ HSPCs. **A.** Schematic comparing *ASXL1* knockout with sgRNAs in exon 2 of the gene to *ASXL1* truncation with sgRNAs in exon 12 near the canonical hotspot. **B.** Edited cells from **A** were grown in liquid culture under multi-lineage differentiation conditions for 7 days, followed by FACS for CD34**. C.** Edited cells were grown in methylcellulose for 14 days and then counted by a blinded observer. Cells were replated and cultured for an additional 14 days and recounted. **D**. Representative images from **C**. **E.** FACS was used to measure engraftment in NSGS mice (hCD45) and myeloid (CD33) vs lymphoid (CD3+CD19) output. **F**. CD34+ cord blood cells expressing mutant *ASXL1* or edited at the *AAVS1* locus (as in Figure 2) were cultured under multi-lineage differentiation conditions for 4 days, followed by RNA-seq (n=3/group). **G**. Colony-forming assay after *FOS* knockout by CRISPR. **H**. Gene set overrepresentation analysis for mutant *ASXL1* up and down gene sets using the transcription factor perturbation expression gene set collection (EnrichR). **I**. Gene set overrepresentation analysis of mutant *ASXL1* up genes using DRUG-seqr. *=p<0.05, **=p<0.01, ***=p<0.001, ****= p<0.0001 as measured by a Student’s T-Test or one-way ANOVA with a Holm-Sidak post-test.

Prior work has shown that mutant ASXL1 requires the BAP1 deubiquitinase for its oncogenic function; however, this has not been functionally validated in human HSPCs^10^. Therefore, we deleted BAP1 in *ASXL1*-mutant and *AAVS1*-control cells and evaluated the impact on HSPC function. BAP1 knockout reversed the effect of mutant ASXL1 on CD34 retention while having a minimal impact on CD34 abundance in control cells (**Supplementary Figure 1B-D**). Further, deletion of BAP1 prevented mutant ASXL1 from increasing replating capacity in the colony-forming assay. We confirmed these findings using a BAP1 inhibitor, which also selectively reversed CD34 retention in *ASXL1*-mutant cells (**Supplementary Figure 1E-G**)^10^.

### Mutant ASXL1 Drives Gene Activation and Repression

**—**To understand the transcriptional changes that enable *ASXL1* mutant cells to resist differentiation, we performed RNA-seq on *ASXL1*-mutant CD34+ HPSCs after four days of culture under pro-differentiation conditions (**Figure 2F**). This revealed markedly decreased expression of the genes *FOS* and *FOSB*, which were downregulated 12 and 7-fold, respectively. These data are the first to support a repressive role for mutant ASXL1 in human hematopoiesis. FOS and FOSB interact with JUN to form the heterodimeric transcription factor AP-1. The role of FOS/AP-1 in hematopoiesis is complex, with some studies showing that high levels of FOS enforce HSC quiescence. In contrast, others suggest that FOS drives HSC activation and differentiation in response to stressors such as inflammation^17–22^. To evaluate whether decreased *FOS* expression reproduces the phenotype of mutant ASXL1, we deleted *FOS* in CD34+ HSPCs. *FOS* knockout increased colony formation in both the first and second rounds of plating, similar to *ASXL1* truncation (**Figure 2G**).

To nominate regulatory pathways responsible for mutant ASXL1-driven transcriptional changes, we performed a gene set overrepresentation analysis using a transcription factor perturbation library. This revealed that mutant ASXL1 upregulates a MECOM gene expression signature and downregulates genes suppressed by CEBPA, among other enriched gene sets (**Figure 2H**). MECOM is a key regulator of long-term hematopoietic stem cell function, whereas CEBPA regulates myeloid differentiation. Recently, CEBPA was identified as a key repressed target downstream of oncogenic MECOM^23,24^. Collectively, these transcriptional changes correlate with the decreased differentiation and increased self-renewal exhibited by *ASXL1*-mutant HSPCs. We also evaluated overlap with the DRUG-seqr database^25^. This revealed a strong enrichment between genes upregulated by mutant ASXL1 and repressed by BET (bromodomain and extra-terminal domain proteins; bind to acetylated histones), or histone deacetylase (HDAC) inhibition (**Figure 2I**). These data implicate aberrant histone acetylation and the dysregulation of core myeloid transcription factors in the increased stemness observed in *ASXL1*-mutant cells.

### Truncating Mutations in ASXL1 Increase Protein Stability

**—**The exact effect of protein truncation on ASXL1 function is not entirely understood, with reports of both protein stabilization and neomorphic function. To evaluate the former in our endogenous model, we introduced HiBiT tags at the ASXL1 locus, one at the mutant and one at the wild-type stop codon (**Figure 3A**)^26^. Despite similar levels of recombination, a significantly higher lytic signal was observed in the *ASXL1*-mutant as compared to the wild-type condition (**Figure 3B** and **Supplementary Figure 2A and 2B**). To confirm that the HiBiT-tagged ASXL1 mutant, we generated single-cell clones from U937 cells and detected a band at the expected molecular weight (**Supplementary Figure 2C**).

**Figure 3.**
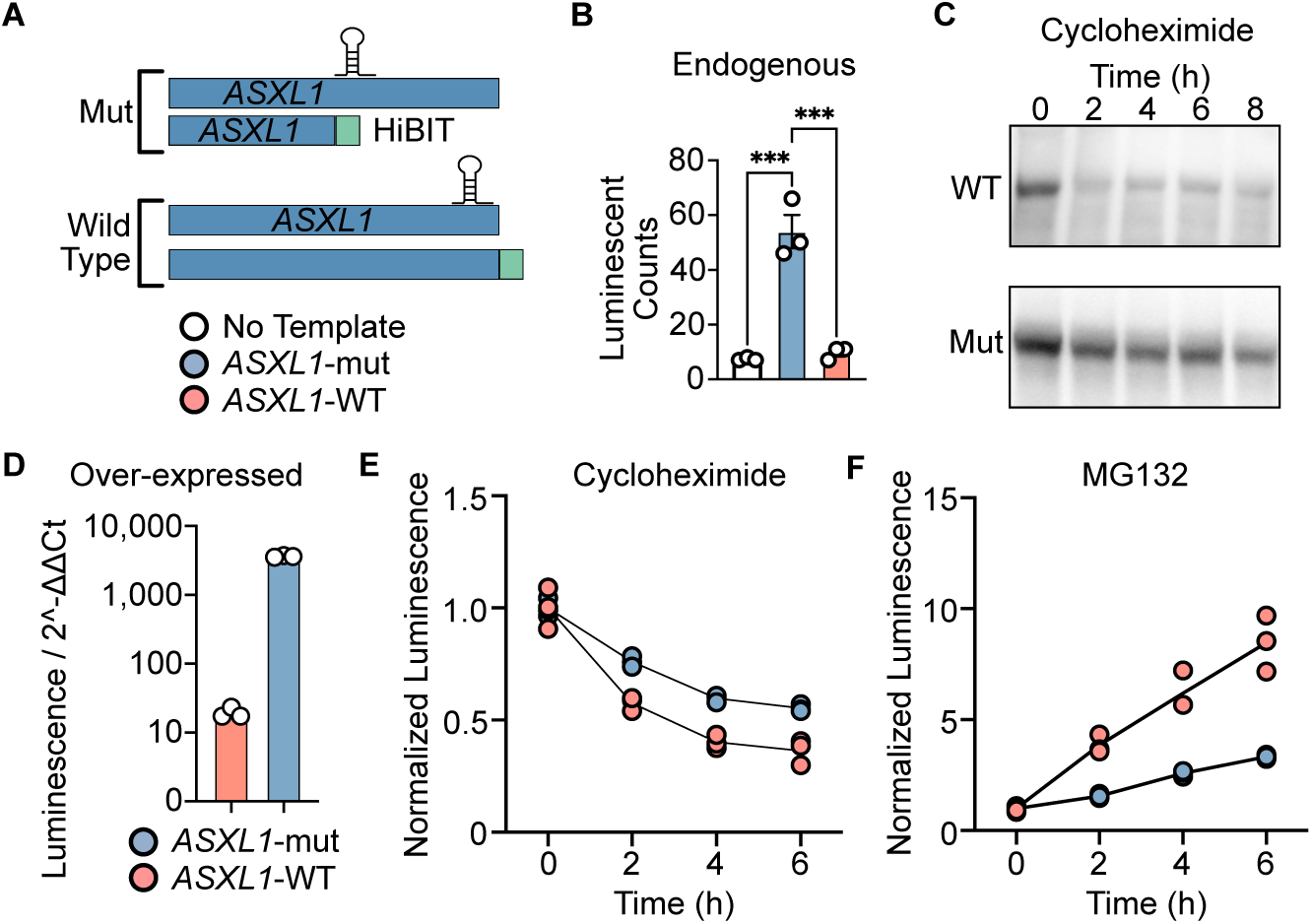
Truncating Mutations in ASXL1 Increase Protein Stability. **A.** Schematic of HiBIT tag knock-in at C632 or the native stop codon of *ASXL1*. **B.** Protein quantification of HiBiT-tagged endogenous ASXL1 by HiBiT lytic assay in CD34+ HSPCs described in A (***=p<0.001). **C.** Anti-FLAG western blot of HEK cells transiently transfected with constructs coding for FLAG-tagged WT or mutant ASXL1 treated with 20 μg/mL cycloheximide. **D.** HiBiT-tagged transiently transfected ASXL1 by HiBiT lytic assay in HEK cells. Luminescent signal normalized to backbone abundance using qPCR, controlling for minor differences in transfection efficiency. **E.** Change in protein levels in HEK cells transiently transfected with HiBiT-tagged ASXL1 by HiBiT lytic assay after treatment with 20 μg/mL cycloheximide (n=3 independent transfections). Signal is normalized to the zero-hour time point to demonstrate relative changes. **F.** Change in protein levels in HEK cells transiently transfected with HiBiT-tagged ASXL1 by HiBiT lytic assay after treatment with 5 μM/mL MG132 (n=3 independent transfections). Signal is normalized to the zero-hour time point to demonstrate relative changes. ***=p<0.001 as measured by a one-way ANOVA with a Holm-Sidak post-test.

To confirm our findings regarding ASXL1 protein stability, we expressed flag-tagged wild-type or mutant ASXL1 in HEK293 cells and performed a cycloheximide time course (**Figure 3C**; **Supplementary Figure 2D**). Wild-type protein appeared to degrade more rapidly than the mutant; however, this interpretation is confounded by the low level of wild-type ASXL1 expression and the marked differences in molecular weight. To control for these confounders, we expressed HiBiT-tagged mutant and wild-type ASXL1 in HEK293 cells and performed a lytic luminescence assay (**Figure 3D**). To control for potential differences in transfection efficiency, we performed qPCR for the GFP backbone and normalized the lytic signal to the relative abundance of GFP. Both the raw and normalized lytic signal for mutant ASXL1 were multiple orders of magnitude higher than that of wild-type ASXL1. To confirm an effect on protein stability, we treated HEK293 cells expressing mutant or wild-type ASXL1 with cycloheximide and monitored the loss of HiBiT lytic signal over time (**Figure 3E**). This revealed a more rapid loss of the wild-type ASXL1 lytic signal than that of mutant ASXL1, suggesting that mutant ASXL1 is more stable. Finally, we used the same system to evaluate the contribution of proteasomal degradation to ASXL1 protein stability. Treatment with the proteasome inhibitor MG132 resulted in a much more rapid accumulation of wild-type ASXL1 than of mutant ASXL1, consistent with altered proteasomal degradation rates (**Figure 3F**). These results are consistent with ChIP-seq data from mutant and wild-type ASXL1, which show similar binding sites but higher binding intensity for mutant ASXL1 (**Figure 1A**)^10^.

### Mutant ASXL1 Interacts with RNA Polymerase II Complex Members

**—**To identify the protein interactions that govern the transcriptional effects of mutant *ASXL1*, we performed a proximity-dependent biotin ligation (BioID) screen using both mutant and wild-type ASXL1 (**Figure 4A**). This identified associations between ASXL1 and multiple components of the basal transcriptional apparatus and RNAPII elongation. We confirmed the previously reported interaction between mutant ASXL1 and BRD4 and found that this also occurs with the wild-type form of the protein. This result corroborates previous proximity-dependent ligation studies of ASXL1 and highlights the utility of this method to capture interactions with WT ASXL1, which often has low protein expression^14^.

**Figure 4.**
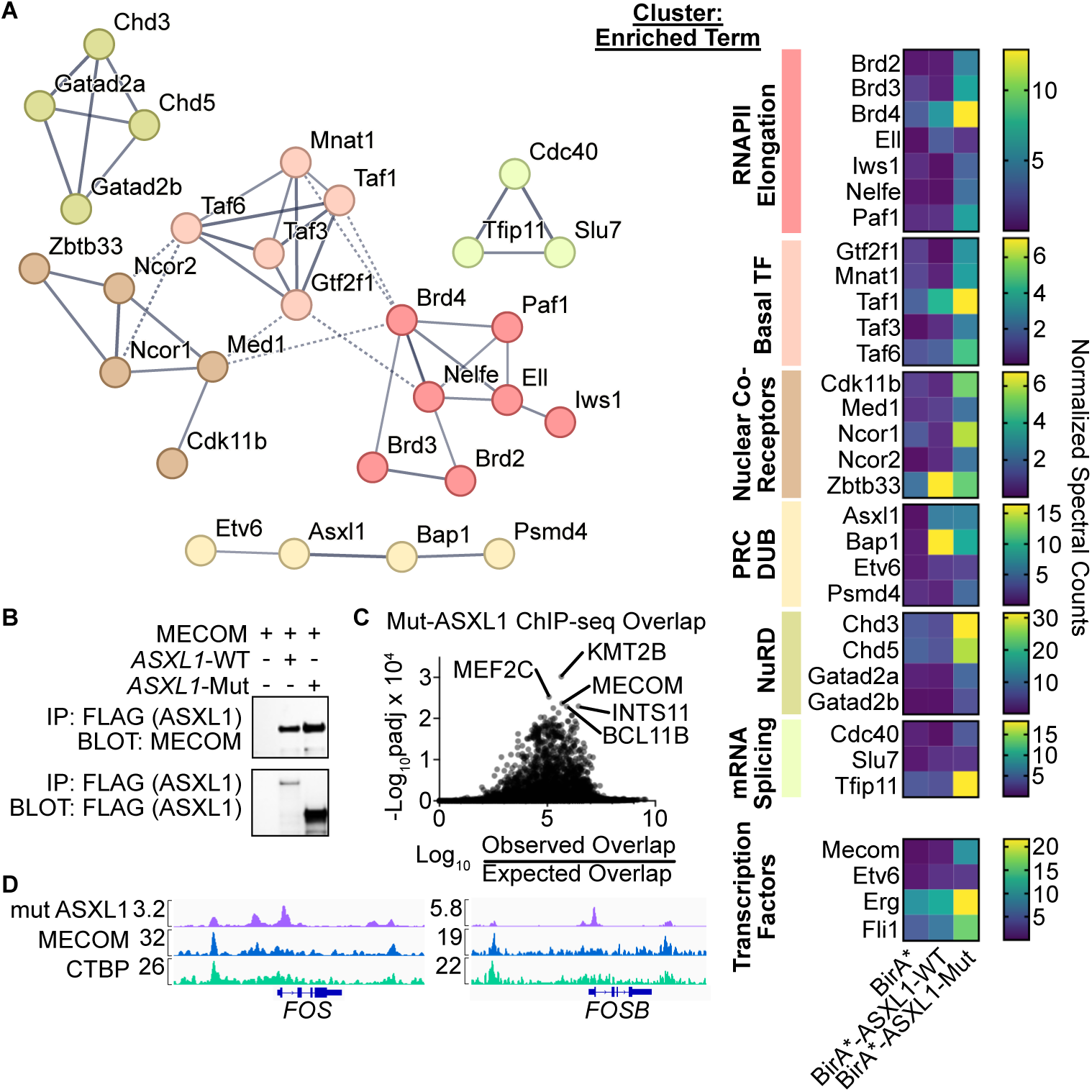
Proximity-Dependent Biotin Ligation for ASXL1. **A.** BioID data identifying an interaction between ASXL1 and RNAPII regulators. Enriched terms identified by STRING/MCL clustering. E=BirA empty vector, W=ASXL1-WT, M=ASXL1-635fs*15. **B.** Immunoprecipitation in HEK293 cells between FLAG-ASXL1 and MECOM. Note that the low expression level of WT ASXL1 is consistent with the known stabilizing effects of mutations. **C.** Intersection of mutant ASXL1 ChIP-seq from Figure 1 with the ReMap 2020 database. **D.** Representative ChIP-seq tracks for ASXL1, MECOM, and CtBP at *FOS* and *FOSB*.

In addition, we identified a previously unappreciated interaction with MECOM. MECOM is recurrently overexpressed in AML via chromosomal translocations and inversions, driving an HSC-like phenotype with increased self-renewal and treatment resistance^15^. It is increasingly appreciated that these oncogenic functions are driven via interaction with CtBP1/2 and transcriptional repression^16^. We confirmed the interaction between MECOM and ASXL1 (both mutant and wild type) using conventional co-immunoprecipitation (**Figure 4B**). We also performed a transcription factor binding site overlap analysis using mutant ASXL1 ChIP-seq data, which revealed MECOM as one of the most highly intersecting factors, including co-binding at mutant ASXL1-repressed genes such as FOS and FOSB (**Figures 4C and 4D**).

### Mutant ASXL1 Interacts with BRD4 to Increase RNAPII Pause Release

**—**To functionally validate these findings, we performed an arrayed CRISPR screen in *ASXL1* mutant CD34+ cells using CD34+ retention as a readout (**Figure 5A**). This revealed that deletion of numerous mutant ASXL1-interacting proteins selectively reduced CD34+ abundance in *ASXL1*-mutant cells compared with *AAVS1*-edited control cells. The strongest hit from this screen was BRD4, a known mutant ASXL1-interacting protein (**Figure 5B**)^12^. BRD4 is required for the release of paused RNAPII, providing a plausible link between the ASXL1 mutation and the abnormal transcriptional kinetics observed in ASXL1-mutant cells. We also identified multiple other mutant ASXL1-interacting chromatin regulators, which were required for mutant *ASXL1*-dependent differentiation resistance. These include *JMJD1C* (H3K9 demethylase) and *SSRP1* (component of the FACT complex). The FACT complex is responsible for nucleosome disassembly during the process of RNAPII-dependent transcription and for nucleosome reassembly after transcription, with deletion leading to aberrant RNAPII pausing. H3K9 methylation governs the formation of heterochromatin, and dysregulation of H3K9 methylation and demethylation can perturb hematopoietic differentiation.

**Figure 5.**
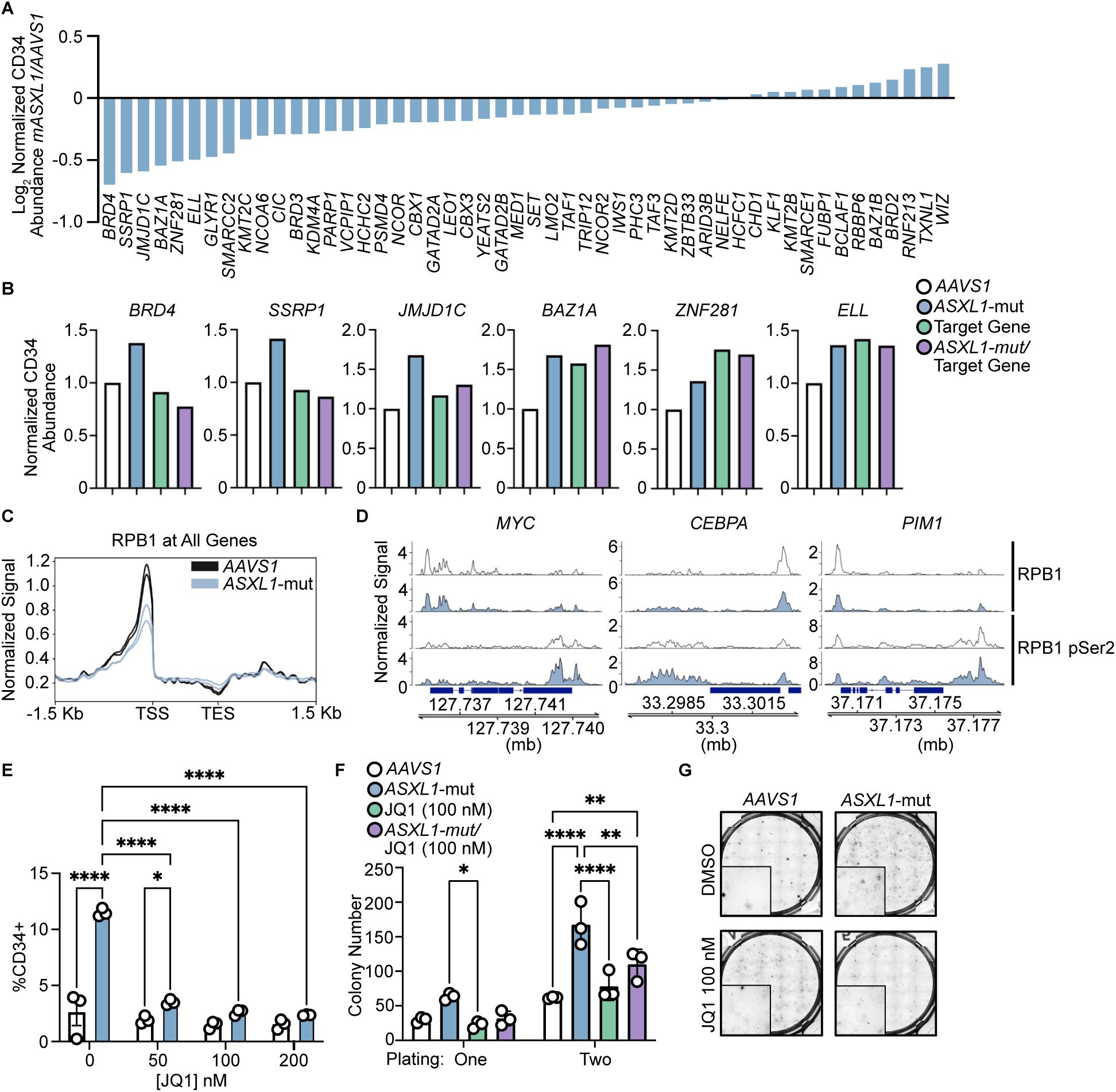
Mutant ASXL1 Leverages BRD4 to Promote Increased Transcription. **A.** Quantification of the impact of target gene knockout on CD34 abundance by first calculating the log2-normalized fold change in CD34 abundance due to knockout of the target gene, then calculating the difference in the log2FC between the AAVS1 and ASXL1-mut cells. **B.** CD34 abundance relative to AAVS1 for top screen hits. **C.** CUT&Tag for RPB1 (large subunit of RNAPII) in *AAVS1* vs *ASXL1*-mut CD34+ cells. **D.** Signal tracks for total and Serine 2 phosphorylated RPB1 in *AAVS1* vs *ASXL1*-mut CD34+ cells. **E.** CD34 abundance by flow cytometry after 7 days of drug treatment in differentiation media (*=p<0.05, ****= p<0.0001). **F.** Colony formation assay with JQ1 treatment. **G.** Representative images from F. *=p<0.05, **=p<0.01, ****= p<0.0001 as measured by two-way ANOVA with a Holm-Sidak post-test.

To understand the functional effects of these protein interactions on transcription in the context of endogenous ASXL1 mutations, we performed CUT&Tag on total and serine 2-phosphorylated RPB1 (large RNAPII subunit) in *ASXL1*-mutant or *AAVS1*-edited CD34+ HSPCs. This revealed globally decreased promoter-bound total RPB1 (paused) and increased TES-bound RPB1 (traveling) (**Figure 5C-D**). Additionally, at certain genes associated with proliferation (i.e., *MYC* and *PIM1*) and hematopoietic cell fate (i.e., *CEBPA*), we observed decreased promoter-bound RPB1 and increased TES-bound serine 2-phosphorylated RPB1, a pattern associated with increased traveling RNAPII. Consistent with this, inhibition of BRD4 with the BET inhibitor JQ1 selectively reduced CD34+ retention in *ASXL1*-mutant cells (**Figure 5E**) and reversed the increased replating capacity of these cells in colony-forming assay (**Figure 5F, G**). Collectively, these results link many oncogenic phenotypes of mutant ASXL1 to increased RNAPII-dependent transcription, in a BRD4-dependent manner.

### Single Cell Transcription Factor Footprinting Identifies Increased MECOM Binding in *ASXL1*-Mutant HSCs

**—**While increased RNAPII-dependent transcription and BRD4 can account for some of the positive effects of mutant ASXL1 on gene expression, they cannot directly explain the strong decrease in immediate early gene expression (**Figure 2F**). MECOM is a likely candidate mediator, given our protein interaction data (**Figures 4A, 4B**), the known repressive role of MECOM^16^, and its established role in governing normal and oncogene-induced HSC self-renewal^15,27^. While low-input techniques for transcription factor profiling (such as CUT&RUN) now permit the interrogation of genome-wide binding within less abundant primary cell types, for the majority of targets, these assays continue to require hundreds of thousands of cells. Given that long-term HSCs (LT-HSCs, the principal site of MECOM action) are rare, we employed a single-cell transcription factor footprinting approach to link MECOM binding to mutant *ASXL1.* We performed sc-ATAC-seq in CD34+ HSCs expressing truncated ASXL1 after three days of culture under differentiation conditions (**Figure 6A**). We then evaluated whether *ASXL1*-mutant HSCs showed changes in MECOM binding. We examined transcription factor footprints at MECOM motifs intersecting with MECOM and ASXL1 ChIP-seq peaks and identified a subset of regions with ASXL1-mutant-specific MECOM footprints, suggesting binding only in *ASXL1*-mutant HSCs (**Figure 6B**)^10,16,28^. These regions showed a clear difference in read pileup between *ASXL1*-mutant and wild-type HSCs within the area centered on the MECOM motif (**Figure 6C**). Gene Ontology analysis revealed a strong enrichment for cell activation pathways, consistent with a role for MECOM in regulating the HSC activation state (**Figure 6D**). To functionally validate these findings, we deleted MECOM in *ASXL1* mutant cells. We found that *MECOM* knockout attenuated the mutant ASXL1-driven resistance to differentiation in liquid culture (**Figure 6E**) and reduced the replating capacity of *ASXL1* mutant cells while only minimally impacting the replating of wild-type cells (**Figure 6F, G**).

**Figure 6.**
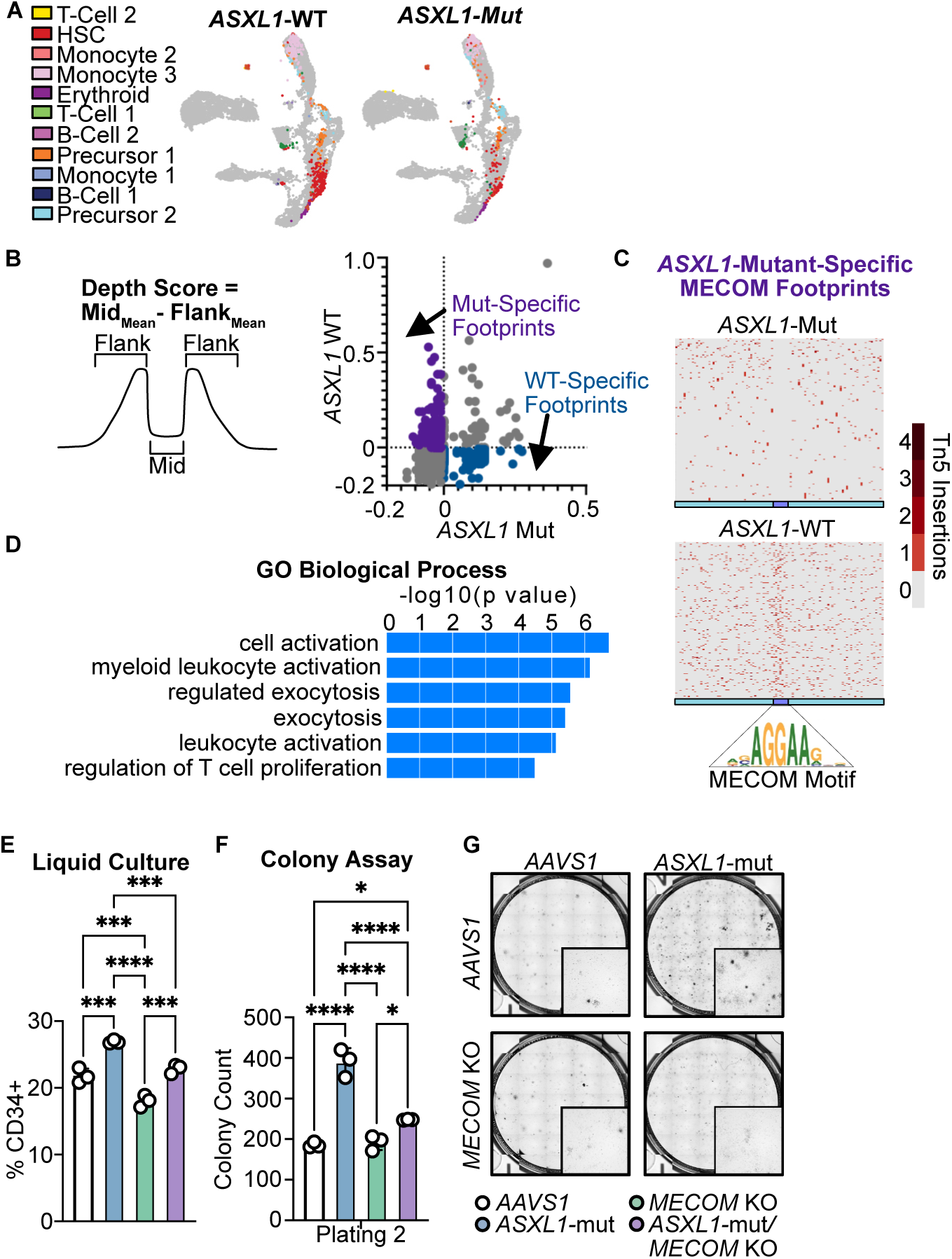
Transcription Factor Footprinting Reveals Increased MECOM Binding in *ASXL1*-Mutant HSCs. **A.** scATAC-seq was performed on CD34+ cord blood cells after 3 days of culture in differentiation medium and was mapped to a healthy bone marrow reference dataset. **B.** Footprint depth scores for TF footprints at MECOM binding sites (from ChIP-seq) that overlap with ASXL1 binding sites (ASXL1-bound, n=1584 sites) in HSCs. Mutant-specific footprints were identified as having a depth score <0 in *ASXL1*-Mut cells and >1 in *ASXL1*-WT cells. **C.** Single locus *ASXL1*-mutant specific MECOM footprints are shown in *ASXL1*-mutant and *ASXL1* wild-type cells (n=249 sites) defined based on differential footprint depth score. **D.** GO analysis was performed on MECOM footprints specific to the *ASXL1*-mutant condition using GREAT. **E.** *ASXL1* mutant or *AAVS1*-edited cells with or without MECOM co-deletion were cultured in multi-lineage differentiation media for 7 days, and CD34 abundance was measured using FACS. **F.** *ASXL1* mutant or *AAVS1*-edited cells with or without MECOM knockout were plated in methylcellulose and counted by a blinded observer after 14 days. No difference in growth between conditions was observed after the initial plating, as in Figure 2. Cells were replated and counted again after 14 days. **G**. Representative Images from **F.** *=p<0.05, ***=p<0.001, ****= p<0.0001 as measured by a two-way ANOVA with a Holm-Sidak post-test.

### Increased MECOM Activity is a Feature of ASXL1-Mutant Myeloid Malignancies

To establish the relevance of MECOM activation in ASXL1-mutant myeloid malignancies, we examined whether *ASXL1*-mutant AML has increased MECOM activity. To do this, we calculated TF activity scores from bulk RNA-seq data for the 630 samples in the Beat AML dataset using the Priori algorithm (**Figure 7A**)^29,30^. Priori leverages known interactions from the Pathway Commons database to predict transcription factor activity. We have previously used Priori to identify novel mediators of drug activity and resistance in AML^29,31^. Differential TF activity score analysis revealed significantly higher MECOM activity in *ASXL1*-mutant AML compared with cases lacking this mutation (**Figure 7B, C**). MECOM gene expression was similar between *ASXL1*-mutant and wild-type cases, demonstrating that the difference in predicted activity is driven by target gene expression (**Supplementary Figure 3A**). None of our *ASXL1*-mutant AML samples had evidence of translocations/inversions involving chromosome 3 or evidence of *MECOM* rearrangement as a potential cause of increased MECOM activity. This suggests that, even in the complex co-mutational landscape of AML, MECOM dysregulation is a hallmark of mutant *ASXL1*.

**Figure 7.**
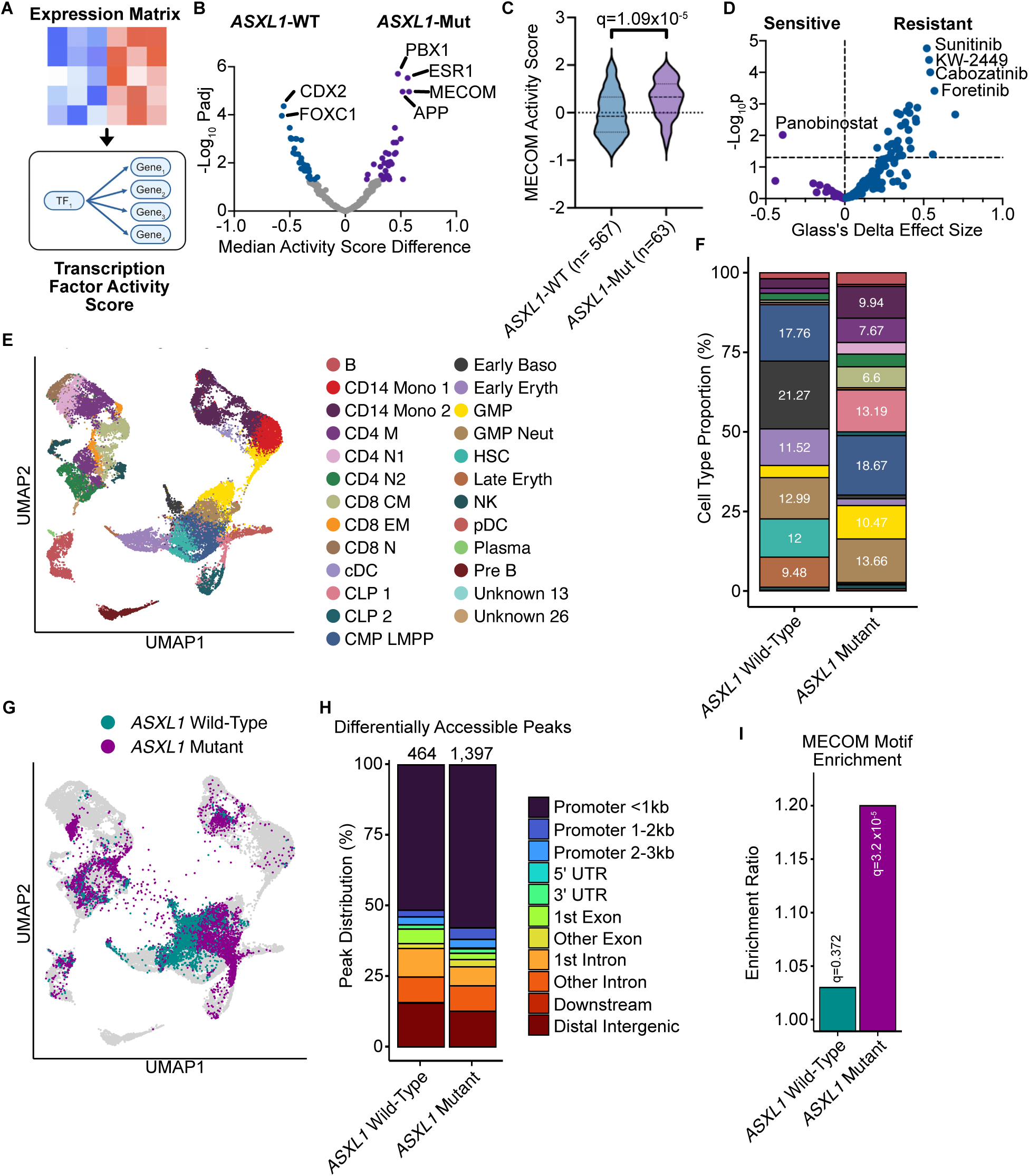
Regulon Analysis Shows Increased MECOM Activity in *ASXL1*-Mutant AML. **A.** Overview of the Priori regulon analysis algorithm, which leverages interactions from the Pathway Commons Database to predict transcription factor activity using expression data. **B.** Differential TF activity score analysis in *ASXL1*-mutant (n=63) and wild-type AML (n=567) using the Beat AML Cohort. **C.** Distribution of MECOM activity scores in *ASXL1*-mutant and *ASXL1*-WT AML samples. None of the *ASXL1*-mutant samples had rearrangements of the MECOM locus. MECOM mRNA counts showed a similar distribution. **D.** Differential drug sensitivity analysis between *ASXL1*-mutant and wild-type AML using the Beat AML cohort. **E.** Reference healthy bone marrow scATAC-seq UMAP with cell type annotations. **F.** Cell type proportions in *ASXL1*-mutant and *ASXL1*-wild type cases (n=2/group). **G.** *ASXL1*-mutant and wild-type AML cells projected onto the reference UMAP. **H.** Peak annotation for differentially accessible regions in *ASXL1* mutant and wild-type cells in all stem and progenitor cells (HSC, CMP-LMPP, GMP, GMP-Neu). **I.** MECOM motif enrichment in differentially accessible peaks from **H.**

To further characterize the epigenetic consequences of mutant *ASXL1* in AML at single-cell resolution, we performed single-cell ATAC-seq (sc ATAC-seq) on n= 4 samples from the Beat AML repository. We used a reference-mapping approach to identify cell types (**Figure 7E**) and observed distinct cell-type proportions between *ASXL1*-mutant and wild-type cases (**Figures 7F and 7G**). We then performed a differential accessibility analysis between *ASXL1*-mutant and wild-type cells within cell type groups with sufficient cell counts in both conditions (**Figure 7H** and **Supplementary Figure 3B**). A database-guided analysis revealed a strong enrichment of SP1 and KLF family member motifs in regions with increased accessibility in *ASXL1*-mutant cells (**Supplementary Figure 3C-F**). SP1 family members bind to CG motifs at the promoters of genes lacking TATA boxes and interact with the basal transcriptional apparatus (i.e. TAFs, also interact with mutant ASXL1 **Figure 4A**) to drive transcriptional initiation^32^. We also evaluated MECOM motif enrichment in our *ASXL1* mutant AML samples. This revealed a significant enrichment of MECOM motifs within differentially accessible peaks in ASXL1 mutant AML cells (**Figure 7I, Supplementary Figure 3G**). Collectively, these data establish that *ASXL1*-mutant leukemias exhibit increased MECOM activity and global transcriptional dysregulation, confirming our findings in ASXL1-mutant HSPCs.

## Discussion

Across the spectrum of myeloid disease, mutations in ASXL1 are associated with high rates of progression, treatment resistance, and increased mortality^2–4^. Developing new treatment approaches that specifically target ASXL1-mutant cells is a significant unmet clinical need. However, a major barrier to this has been the lack of model systems in which mutant ASXL1 alone produces oncogenic phenotypes. In this work, we leveraged a CRISPR-engineered CD34+ HSPC system and protein-protein interaction data to identify novel ASXL1-specific dependencies. We find that mutant ASXL1 interacts with both gene-activating and repressive proteins in HSPCs, both of which are required for the complete oncogenic phenotype. A BRD4/RNAPII-dependent mechanism governs mutant ASXL1-dependent transcriptional activation, whereas a previously unappreciated interaction with the transcription factor MECOM mediates gene repression. Both pathways are amenable to therapeutic inhibition, providing a translational path for the developing mutant ASXL1-specific therapy.

A key finding of our work is that truncating mutations in ASXL1 increase protein stability, leading to protein level overexpression. In contrast to prior studies, our BioID experiments did not detect any neomorphic protein interactions with mutant ASXL1^12^. Consistent with this, mutant ASXL1 ChIP-seq data show near-identical localization but increased abundance compared to wild-type ASXL1. This argues that ASXL1 mutations primary function through protein-level overexpression and increased activity of ASXL1-containing chromatin complexes. Indeed, BAP1 is known to deubiquitinate and stabilize ASXL1^13^. This provides a plausible explanation for the requirement for BAP1 in mutant ASXL1-dependent oncogenesis, despite ASXL1 and H2aK119Ub showing mutually exclusive genome-wide binding profiles. Our data further argues that a significant function of the C-terminus of ASXL1 is to destabilize the protein. The specific motifs required for this function and the process through which it is degraded represent an exciting avenue for future investigation.

Prior studies identified BRD4 as a mutant ASXL1-interacting protein and found that mutant *Asxl1*-expressing mouse bone marrow cells exhibit increased cell death in response to BET inhibitor treatment^12^. However, the mechanism by which BRD4 contributes to mutant ASXL1-driven transcription and oncogenic phenotypes had not previously been described. Our work here demonstrates that *ASXL1*-mutant HSPCs exhibit increased RNA polymerase II elongation, consistent with increased BRD4-dependent RNAPII pause-release. Furthermore, chemical or genetic inhibition of BRD4 selectively differentiates *ASXL1*-mutant cells and reduces their clonogenicity in a colony-forming assay. These findings functionally support the requirement for BRD4 in driving at least a component of mutant ASXL1-driven oncogenesis and provide strong pre-clinical rationale for leveraging drugs targeting BET proteins for the treatment of *ASXL1*-mutant myeloid malignancies.

While prior studies have attempted to define the protein interactome of mutant ASXL1, these have been done with lower-sensitivity immunoprecipitation mass spec approaches in non-hematopoietic cell types. Therefore, it is unsurprising that our Bio-ID experiments identified numerous new context-specific interacting proteins, including MECOM. Our findings indicate that mutant ASXL1 is overexpressed and thus may recruit more MECOM to a subset of target genes, driving gene repression and increased stemness. MECOM activity is increased in *ASXL1*-mutant myeloid malignancies, and ASXL1 mutation in *MECOM*-rearranged AML further blocks differentiation, supporting mechanistic interdependence. These findings are paradigm-shifting and underscore the importance of studying global epigenetic regulators in their appropriate cellular context.

Our results point to multiple potential avenues for therapeutically targeting mutant ASXL1 chromatin complexes, a significant unmet clinical need. BET inhibitors are already well advanced in clinical development and have favorable safety and tolerability profiles^33,34^. These drugs could easily be explicitly repurposed for *ASXL1*-mutant myeloid malignancies. CDK9 inhibition is also a promising approach for inhibiting aberrant RNAPII pause-release in ASXL1 mutant cells. Recently, a 50% response rate in relapsed/refractory *ASXL1*-mutant AML was observed using the CDK9 inhibitor tambiciclib (SLS-009), providing the impetus for the initiation of front-line clinical trials^35^. If these trials confirm the initial promise of this approach, tambiciclib could become the standard-of-care for *ASXL1*-mutant myeloid malignancies. Finally, MECOM also represents a potential approach to targeting ASXL1-mutant myeloid malignancies. Recent work has demonstrated that MECOM-dependent oncogenesis depends on interaction with CTBP1/2 co-repressors. Peptide-based inhibitors of the MECOM-CTBP protein interface selectively reverse MECOM-induced gene repression, driving AML cell differentiation^16^. It remains to be established whether therapeutic targeting of the MECOM-CTBP interface will have a sufficient therapeutic index to avoid toxicity to healthy stem cells.

Collectively, our results provide new mechanistic insights into the mechanisms of dual activating and repressive functions of mutant ASXL1 and highlight new therapeutic opportunities for a difficult-to-treat subset of myeloid malignancies.

## Methods

### CD34 Cell Culture

Healthy human cord blood was obtained from Northwest Bloodworks. Mononuclear cells were isolated by Ficoll-paque density gradient centrifugation and ACK lysis. CD34+ cells were isolated from mononuclear cells using MACS, following the manufacturer’s protocol (Miltenyi Biotec, CD34 MicroBead Kit, human, Cat. #130-046-702). After isolation, CD34+ cells were frozen in freezing medium (90% FBS, 10% DMSO). CD34+ cells were cultured in expansion media (StemSpan SFEM II (StemCell Technologies Cat. #09605) supplemented with 10% StemSpan CD34+ Expansion Supplement (StemCell Technologies Cat. #02691) and 0.2% UM729 (StemCell Technologies Cat. #72332).

For differentiation assays, CD34+ cells were transferred to differentiation media (StemSpan SFEM II (StemCell Technologies Cat. #09605) supplemented with 100 ng/mL SCF (PeproTech, Cat. #300-07), 100 ng/mL FLT3L (PeproTech, Cat. #300-19), 100 ng/mL IL6 (PeproTech, Cat. #200-06), 20 ng/mL IL3 (PeproTech, Cat. #200-03), 100 ng/mL TPO (PeproTech, Cat. #300-18), 2 ng/mL EPO (PeproTech, Cat. #100-64), 10 ng/mL GM-CSF (PeproTech, Cat. #300-03)) immediately after CRISPR editing. After incubating for seven days, the cells (300,000) were stained in 100 uL of staining buffer (BD Biosciences, Cat. #554656) with 2 uL of BV421 anti-human CD34 antibody (BioLegend Cat. #343610). Cells were analyzed on a BD FACSymphony A5 flow cytometer.

### Colony assays

After CRISPR editing, cells were grown in expansion media for 24-48 hours. 5 x 10^5^ to 1 x 10^6^ cells were then stained in 100 µL of staining buffer (BD Biosciences, Cat. #554656) with 5 µL of BV421 anti-human CD34 antibody (BioLegend Cat. #343610), 20 µL of PE anti-human CD49c/ITGA3 antibody (BD Biosciences Cat. #556025), 5 µL of APC-Cy7 anti-human CD45RA antibody (BioLegend Cat. #304127), 5 µL of PerCP-Cy5.5 anti-human CD90 antibody (BioLegend Cat. #328118), 3 µL of FITC anti-human EPCR antibody (Miltenyi Biotec Cat. #130-129-886), and 5 µL of APC anti-human CD133 antibody (BioLegend Cat. #393906). 6000 long-term HSCs per condition (CD34+, ITGA3+, EPCR+, CD90+, CD45RA-) were sorted by FACS directly into 4 mL of MethoCult H4434 (StemCell Technologies Cat. #04434) supplemented with 1% Penicillin-Streptomycin and 0.1% Fungizone. Cells were mixed by vortexing, and 1 mL of media and cells was plated in triplicate in a 6-well SmartDish (StemCell Technologies Cat. #27370). After incubating for 14 days, plates were imaged on a STEMvision (StemCell Technologies). For replating, cells were washed with IMDM supplemented with 20% FBS before plating 100,000 cells split into triplicate. Colonies were counted manually in ImageJ after being blinded by another investigator.

### CRISPR Editing

CD34+ cord blood cells were grown in expansion media for 5-7 days before nucleofection. Nucleofection was performed using a Lonza P3 Primary Cell 4-D Nucleofector X Kit S (Cat. # V4XP-3032) in a Lonza 4D-Nucleofector X Unit (Cat. # AAF-1003X). For each reaction, 1.23 ng of sgRNA and 20 pmol of SpCas9 were combined in P3 buffer before incubation for 30 minutes. 1 x 10^6^ cells were then added to each reaction and nucleofected using the DZ-100 program. For the HiBiT knock-in protocol, 100 pmol of HDR template were added to the sgRNA and spCas9 mixture.

### Co-immunoprecipitation

For co-immunoprecipitation studies, pMYSIG_Flag_ASXL1_WT or pMYSIG Flag_ASXL1_MT2 plasmids were co-transfected with pSmal_MECOM (variant 6) into HEK293 cells with Fugene 6 (Promega Cat. #E2651) following the manufacturer’s protocol. Cells were harvested 48 hours later with Pierce IP lysis buffer (PI87788) plus protease inhibitor cocktail (Roche Cat. #1169748001) and 1 mM phenylmethanesulfonylfluoride solution (Sigma Cat. #93482). 1 mg of total cell lysate was incubated with 2 µg anti-FLAG antibody (Sigma Cat. #F1804) overnight at 4 °C. The lysates were incubated with 40 μL prewashed protein A/G plus-agarose (Santa Cruz Biotechnology Cat. #sc-2003) for 3 hours at 4°C. Beads were washed with cold IP buffer 4 times to remove unbound proteins and centrifuged at 2500 rpm for 5 minutes at 4 °C. The samples were eluted with 1X GS buffer (2% sodium dodecyl sulfate, 50mM Tris, pH 8, 10% glycerol, and 0.03% bromophenol blue) and prepared for SDS-PAGE. SuperSignal West Pico PLUS Chemiluminescent (ThermoFisher Cat. #34580) was used for detection on an iBright 1500 Imager (ThermoFisher).

### Proximity Dependent Biotin Ligation

20 million cells were harvested after treating for 24 hours with 50 uM biotin and 1 ug/ml doxycycline to induce expression of BirA-ASXL1 fusion proteins. Cells were then lysed in 540 µL lysis buffer (8M urea in 50 mM Tris pH 7.4 with 1x cOmplete protease inhibitor (Roche Cat. #11697498001)) before adding 25 U Pierce Universal Nuclease (ThermoFisher Cat #88700). After 10 minutes of incubation, Triton-X 100 was added to a final concentration of 1%. Samples were then sonicated on a QSonica sonicator at 30% amplitude with a pulse of 3 seconds on, 7 seconds off, for a total of 30 seconds on (10 cycles). Samples then rested on ice for 1 minute before sonication was repeated for a total of three rounds. After sonication, 1260 µL lysis buffer was added before centrifugation for 10 minutes at 16,500g and 4°C. 1 mL of supernatant was then mixed with 200 µL of washed gelatin sepharose beads before incubating for 2 hours on a rotator at 4°C. Samples were then centrifuged for 5 minutes at 800g and 4°C. The supernatant was mixed with 50 µL washed streptavidin sepharose beads before incubating for 4 hours at 4°C. Samples were then centrifuged for 5 minutes at 800g and 4°C before removing the supernatant. The pellets were washed 5x with 1 mL wash buffer (50 mM Tris pH 7.4 with 8M urea) and washed once with 1 mL ammonium bicarbonate buffer (50 mM ammonium bicarbonate in 50 mM Tris pH 7.4) before resuspending in 50 mL ammonium bicarbonate biotin buffer (50 mM ammonium bicarbonate and 1 mM biotin in 50 mM Tris pH 7.4). Samples were then flash frozen and stored at −80°C prior to mass spectrometry.

### Bulk RNA-seq

After CRISPR editing, CD34+ HSPCs were grown in differentiation media for four days. CD34+ cells were then re-isolated using MACS (Miltenyi Biotec, CD34 MicroBead Kit, human, Cat. #130-046-702) and following the manufacturer’s protocol. CD34+ cells were pelleted by centrifugation and resuspended in 350 µL RLT (Qiagen Cat. # 79216). Qiashredder (Qiagen Cat. #79656) columns were used to homogenize the cells before proceeding with RNA extraction using an RNeasy Micro Kit (Qiagen Cat. #74004) following the manufacturer’s protocol. RNA was quantified, and integrity was verified on a TapeStation using an RNA Kit (Agilent Cat. # 5067-5576). Library preparation was performed using the Watchmaker mRNA Library Prep Kit (Cat. #7BK0001) with 350 ng of input RNA. Libraries were quantified using a TapeStation HS D1000 Kit (Agilent Cat. #5067-5584), pooled, and sequenced on an Illumina NextSeq 2000 with 61 bp paired-end reads.

### S3-ATAC-seq

Samples were prepared for s3-ATAC seq as previously described^36^. Briefly, 96 uniquely indexed transposome complexes were assembled using previously described methods^37^. Complexes were diluted to 2.5 µM in a protein storage buffer composed of 50% (v/v) glycerol (Sigma G5516), 100 mM NaCl (Fisher Scientific S271-3), 50 mM Tris pH 7.5 (Life technologies AM9855), 0.1 mM EDTA (Fisher Scientific AM9260G), 1 mM DTT (VWR 97061-340) and stored at −20 °C. At the time of nuclei dissociation, 50 ml of nuclei isolation buffer (NIB-HEPES) was freshly prepared with final concentrations of 10 mM HEPES-KOH (Fisher Scientific, BP310-500 and Sigma Aldrich 1050121000, respectively), pH 7.2, 10 mM NaCl, 3 mM MgCl2 (Fisher Scientific AC223210010), 0.1% (v/v) IGEPAL CA-630 (Sigma Aldrich I3021), 0.1% (v/v) Tween (Sigma Aldrich P-7949) and diluted in PCR-grade Ultrapure distilled water (Thermo Fisher Scientific 10977015). After dilution, two tablets of Pierce Protease Inhibitor Mini Tablets, EDTA-free (Thermo Fisher, A32955) were dissolved and suspended to prevent protease degradation during nuclei isolation. Tagmentation plates were prepared by the combination of 420 µl of 1,400 nuclei per µl of solution with 540 µl 2× TD Buffer (Nextera XT Kit, Illumina Inc. FC-131-1024). From this mixture, 8 µl (roughly 5,000 nuclei in total) was pipetted into each well of a 96-well plate dependent on well schema (Fig. 1b). Then 1 µl of 2.5 µM uniquely indexed transposase was then pipetted into each well. Tagmentation was performed at 55 °C for 10 min on a 300 r.c.f. Eppendorf ThermoMixer. Following this incubation, plate temperature was brought down with a brief incubation on ice to stop the reaction. Dependent on experimental schema pools of tagmented nuclei were combined and 2 µl of 5 mg ml−1 4,6-diamidino-2-phenylindole (DAPI) (Thermo Fisher Scientific D1306) were added.

Nuclei were then flow sorted via a Sony SH800 to remove debris and attain an accurate count per well before PCR. A receptacle 96-well plate was prepared with 9 µl of 1× TD buffer (Nextera XT Kit, Illumina Inc. FC-131-1024,diluted with ultrapure water) and held in a sample chamber kept at 4 °C. Fluorescent nuclei were then flow sorted gating by size, internal complexity and DAPI fluorescence for single nuclei following the same gating strategy as previously described40. Immediately following sorting completion, the plate was sealed and spun down for 5 min at 500 r.c.f. and 4 °C to ensure nuclei were within the buffer.

Nucleosomes and remaining transposases were then denatured with the addition 1 µl of 0.1% SDS (roughly 0.01% final concentration) per well. Then 4 µl of NPM (Nextera XT Kit, Illumina Inc.) per well was subsequently added to perform gap-fill on tagmented gDNA, with an incubation at 72 °C for 10 min. Next, 1.5 µl of 1 µM A14-LNA-ME oligo was then added to supply the template for adapter switching. The polymerase-based adapter switching was then performed with the following conditions: initial denaturation at 98 °C for 30 s, ten cycles of 98 °C for 10 s, 59 °C for 20 s and 72 °C for 10 s. The plate was then held at 10 °C. After adapter switching 1% (v/v) Triton-X 100 in ultrapure H2O (Sigma 93426) was added to quench persisting SDS. At this point, some plates were stored at −20 °C for several weeks while others were immediately processed.

The following was then combined per well for PCR: 16.5 µl of sample, 2.5 µl of indexed i7 primer at 10 µM, 2.5 µl of indexed i5 primer at 10 µM, 3 µl of ultrapure H2O, 25 µl of NEBNext Q5U 2× Master mix (New England Biolabs M0597S) and 0.5 µl of 100× SYBR Green I (Thermo Scientific S7563) for a 50 µl of reaction per well. A real-time PCR was performed on a BioRad CFX with the following conditions, measuring SYBR fluorescence every cycle: 98 °C for 30 s; 16–18 cycles of 98 °C for 10 s, 55 °C for 20 s, 72 °C for 30 s, fluorescent reading, 72 °C for 10 s. After fluorescence passes an exponential growth and begins to inflect, the samples were held at 72 °C for another 30 s then stored at 4 °C.

Amplified libraries were then cleaned by pooling 25 µl per well into a 15-ml conical tube and cleaned via a Qiaquick PCR purification column following the manufacturer’s protocol (Qiagen 28106). The pooled sample was eluted in 50 µl 10 mM Tris-HCl, pH 8.0. Library molecules then went through a size selection via SPRI selection beads (Mag-Bind TotalPure NGS Omega Biotek M1378-01). Next, 50 µl of vortexed and fully suspended room temperature SPRI beads were combined with the 50-µl library (1× clean up) and incubated at room temperature for 5 min. The reaction was then placed on a magnetic rack and once cleared, supernatant was removed. The remaining pellet was rinsed twice with 100 µl of fresh 80% ethanol. After ethanol was pipetted out, the tube was spun down and placed back on the magnetic rack to remove any lingering ethanol. Next, 31 µl of 10 mM Tris-HCl, pH 8.0 were then used to resuspend the beads off the magnetic rack and allowed to incubate for 5 min at room temperature. The tube was again placed on the magnetic rack and once cleared, the full volume of supernatant was moved to a clean tube. DNA was then quantified by Qubit double-stranded DNA High-sensitivity assay following the manufacturer’s instructions (Thermo Fisher Q32851). Libraries were then diluted to 2 ng µl−1 and run on an Agilent Tapestation 4150 D5000 tape (Agilent 5067-5592). Library molecule concentration within the range of 100–1,000 base pairs (bp) was then used for final dilution of the library to 1 nM. Diluted libraries were then sequenced on a NextSeq 2000 (Illumina).

### CUT&Tag

Prior to performing CUT&Tag, CD34+ HSPCs were fixed. Fixative buffer was prepared by diluting a freshly opened vial of 16% Formaldehyde (CST, 12606) to a concentration of 0.1% in serum-free DMEM (Gibco, 10569010). Up to 3 million cells in suspension were transferred into 1.5 ml tubes and pelleted at 300 × g for 7 minutes. Media was removed, and pellets were resuspended in 1 ml of fixative buffer. They were incubated at room temperature for 3 minutes. Cells were pelleted at 300 × g for 7 minutes without quenching. The supernatant was discarded. Fixed cells were washed in 1 ml of DMEM supplemented with 0.1% BSA, pelleted again, and cryopreserved in 10% DMSO/90% Fetal Bovine Serum. Cryopreserved cells were stored at - 70°C for long-term preservation.

Cells were counted, and 360,000 cells per antibody per condition were harvested by centrifugation at 300g. Cells were then washed in 1 mL PBS before resuspending in 1 mL cold NE2B buffer (25 mM HEPES-KOH pH 7.9, 12.5 mM KCl, 0.125% Triton X-100, and 0.5 mM spermidine trihydrochloride (MP Biomedicals Cat. #0210047201) in molecular grade water) and incubating for 10 minutes on ice to extract nuclei. Nuclei were spun down and resuspended at 1 x 106 nuclei per mL in cold NE2B buffer. For each sample, 100 µL of nuclei were mixed by gentle vortexing with 10 µL of ConA beads (Bangs Laboratories) that were washed 3 times with ConA-Binding buffer (20 mM HEPES-KOH pH 7.9, 10 mM KCl, 1 mM CaCl2, and 1 mM MnCl2 in molecular grade water) before incubating for 10 minutes. Samples were then placed on a magnet stand, and the supernatant was removed. Primary antibodies (Mouse ⍺RPB1 CTD (Cell Signaling Technology Cat. #2629), Rabbit ⍺RPB1 Ser2P (Cell Signaling Technology Cat. #13499), Rabbit ⍺H3K27me3 (Cell Signaling Technology Cat. #9733), Rabbit ⍺H2aK119Ub (Cell Signaling Technology Cat. #8240), Rabbit IgG (Cell Signaling Technology Cat. #2729)) were diluted 1:100 in antibody buffer (20 mM HEPES pH 7.5, 150 mM NaCl, 0.5 mM spermidine trihydrochloride, 2 mM EDTA, 1X protease inhibitor (Pierce mini-tablets EDTA-free Cat. #A32955), and 0.01% digitonin in molecular grade water) before being added to respective samples and mixing by gentle vortexing. Samples were then placed on a nutator overnight at 4°C. After incubation, samples were placed on a magnet stand, and the supernatant was removed. Secondary antibodies (Guinea pig anti-rabbit (Antibodies Online Cat. #ABIN101961), Rabbit anti-mouse (Abcam Cat. #ab6709)) were diluted 1:100 in cold Digitonin-150 buffer (20 mM HEPES pH 7.5, 150 mM NaCl, 0.5 mM spermidine trihydrochloride, 1X protease inhibitor, and 0.01% digitonin in molecular grade water) before being added to respective samples and mixing by gentle vortexing. Samples were then placed on a nutator at room temperature for 30 minutes. After incubation, samples were washed twice with 200 µL of cold Digitonin-150 buffer before resuspending in 50 µL of Digitonin-300 buffer (20 mM HEPES pH 7.5, 200 mM NaCl, 0.5 mM spermidine trihydrochloride, 1x protease inhibitor, and 0.01% digitonin in molecular grade water) with 2.5 µL of pA/G-Tn5 (Epicypher Cat. #15-1017) and gently vortexing to mix. Samples were then placed on a nutator at room temperature for 1 hour. After incubation, samples were washed twice with 200 µL cold Digitonin-300 buffer and the supernatant was removed. Samples were resuspended in 50 µL cold Tagmentation buffer (Digitonin-300 buffer supplemented with 10 mM MgCl2) and incubated in a thermocycler for 1 hour at 37°C (lid temperature 47°C). Samples were then placed on a magnet stand, and the supernatant was removed before washing with 50 µL of TAPS Wash buffer (10 mM TAPS pH 8.5 and 0.2 mM EDTA in molecular grade water). After removing the wash buffer, DNA was eluted by adding 5 µL SDS release buffer (0.1% SDS and 10 mM TAPS pH 8.5 in molecular grade water) and vortexing before incubating in a thermocycler at 58°C for 1 hour (lid temperature 68°C). Samples were removed from the thermocycler and 15 µL of Triton neutralization buffer (0.67% Triton-X100 in molecular grade water) was added. Samples were PCR amplified using custom primers at 1 uM and NEBNext High Fidelity 2X PCR mix (New England Biolabs Cat. #M0541) under the following conditions: 5 minutes at 58°C, 5 minutes at 72°C, 45 seconds at 98°C, 16 cycles of 15 seconds at 98°C and 10 seconds at 60°C, and 60 seconds at 72°C. Samples were purified using Omega Bio-Tek Mag-Bind TotalPure NGS beads at 1.3x volume before elution in 10 mM Tris-HCl pH 8.0. Samples were quantified using a Qubit fluorometer (1X dsDNA High Sensitivity Kit (Invitrogen Cat. #Q33230)) and on a TapeStation (HS D1000 Kit) before pooling and sequencing on a NextSeq 2000 with 61 bp paired-end reads.

### Multiplexed single-cell ATAC-seq on Primary AML Patient Samples

Cryopreserved primary AML patient samples were thawed, and dead cells were removed using a Dead Cell Removal Kit (Miltenyi, 130-090-101). Cells were then pre-indexed using a Pre-Indexing Kit for scATAC-seq (Scale Biosciences). Briefly, this involves tagmentation with 24 independently indexed transposomes, followed by pooling and superloading onto a Chromium Controller (10X Genomics) at a targeted cell recovery of 100,000 cells per channel. The Tn5 index enables the assignment of fragments from multiple cells within a single droplet to their respective cells of origin. The remainder of the library preparation was performed according to the standard protocol for the single-cell ATAC-seq V2 workflow (10X Genomics). Libraries were sequenced on a NextSeq 2000 using the following sequencing protocol: R1: 36 cycles; i7: 8 cycles; i5: 16 cycles; R2: 63 cycles.

### RNA-seq analysis

RNA-seq reads were processed using an in-house pipeline (https://github.com/maxsonBraunLab/rna_seq). Reads were trimmed with fastp, then aligned using STAR^38^. A counts table was generated for protein-coding genes, and differentially expressed genes were identified with DESeq2 with an adjusted p-value cutoff of 0.05. Gene set overrepresentation analysis was performed using EnrichR^39^ and SeqR^25^.

### CUT&Tag Analysis

CUT&Tag data was processed using an in-house pipeline (https://github.com/maxsonBraunLab/cutTag-pipeline). Reads were trimmed with FastP^40^ and aligned with Bowtie2^41^. Tracks were generated using deepTools^42^, and replicate tracks were merged using WiggleTools^43^. Track visualizations were made using deepTools and Gviz^44^. Correlation plots were made using deepTools using 100 bp bins.

### ChIP-seq Analysis

Publicly available ChIP-seq datasets were processed using the Nextflow ChIP-seq pipeline^45^. Reads were trimmed using Trim Galore^46^ and aligned using BWA^47^. Duplicates were marked using Picard^48^ and reads were filtered using SAMtools^49^ and BAMtools^50^. BigWig files were generated using BEDTools^51^, and peaks were called using MACS2^52^. Consensus peaks were identified using BEDTools. Genomic region annotation of peaks was performed using ChIPseeker^53^.

### S3 ATAC Analysis and Footprinting

After sequencing, data were converted from BCL format to FASTQ format using bcl2fastq (v.2.19.0, Illumina Inc.) with the following options: with-failed-reads, no-lane-splitting, fastq-compression-level = 9, create-fastq-for-index-reads. Reads were then demultiplexed with the python implementation of unidex (https://github.com/ohsu-cedar-comp-hub/unidex), accounting for the 4-nucleotide insert from the Scale transposase before index 3. Briefly, reads were assigned to their expected primer index sequence allowing for sequencing error (Hamming distance ≤2) and indexes were concatenated to form a cellID. The Tn5 barcodes were then used to parse reads into per-sample FASTQ files. Each sample was aligned to the human genome (GRCh37/hg19) using bwa-mem^54^ with default parameters. Reads aligned to scaffolds, the mitochondrial genome, or Y chromosome were removed, as well as insert sizes below 20 or above 10,000 nucleotides. Deduplication was then performed on BAM files to remove all paired reads aligned to the same coordinates within the same cellID.

ArchR^55^ was used for quality control filtering and clustering, retaining cellIDs with a minimum TSS enrichment of 3 and minimum 1,000 fragments. Dimensionality reduction was performed with addIterativeLSI and the following parameters: iterations = 5, clusterParams = list(resolution = 1.5, sampleCells = 750, maxClusters = 12), varFeatures = 25000, and dimsToUse = 20. Clustering was performed with addClusters as a wrapper for the Seurat^56^ FindClusters function with the following parameters: resolution = 1.5, maxClusters = 12, sampleCells = 750. To assign cell type, query cells were mapped to a healthy hematopoiesis scATAC dataset^57^ reprocessed with Signac^58^ to be used as a reference for the Seurat v.4.3.0 anchor-based reference mapping framework.

Footprinting analysis was performed on *ASXL1*-mutant and wild-type cells annotated as HSC from mapping to the reference dataset. FIMO^59^ was used to detect motif matches for an EVI1-like ETS family motif derived from MECOM regulatory studies genome-wide^27^. Motif matches within MECOM ChIP-seq peaks^16^ and within ASXL1 consensus ChIP-seq peaks between mutant and wild type^10^ were considered. Each of the 1,580 motif matches were extended from the center to reach a total length of 100 base pairs, and footprint depth scores were computed for each flanked motif match in each condition. Briefly, Tn5 insertions (found at fragment ends) were summed across the cells at each base pair, and the depth score was computed as the mean insertions in the motif region minus the mean insertions in the flanks, where a score below zero indicates the presence of a footprint^60^.

### Beat AML Analysis

Analysis of the BeatAML cohort was performed using the raw and analyzed published data from the cohort^30,61^. Priori scores for the cohort were calculated in the prior manuscript^29^ and compared using an FDR-corrected Welch’s t-test. Differential drug responses were calculated from inhibitor area under the curve measurements from the BeatAML manuscript and compared between ASXL1-mutant and wild-type cases using Glass’s delta effect size, as previously reported^30^.

### AML Patient Sample scATAC Analysis

The pre-indexed scATAC-seq libraries were pre-processed and aligned to the human genome (GRCh38/hg38) using a Nextflow pipeline from Scale Biosciences (v1.1, https://github.com/ScaleBio/ScaleTagToolkit). Reads were aligned using Bowtie 2^41^, and deduplicated BAM files were generated with the pipeline’s default settings.

ArchR^55^ was used for cell quality control filtering, retaining cells with transcription start site (TSS) enrichment >= 15 and at least 1,000 unique fragments. Retained cells were analyzed using Seurat^56^ and Signac^58^. Normalization and dimensionality reduction were performed using latent semantic indexing (LSI), which combines term frequency-inverse document frequency (TF-IDF) normalization with singular value decomposition (SVD). Cells were clustered within each sample, and peaks were identified using MACS2^58^. Peaks from samples were merged to generate a single consensus peak set for differential accessibility analysis. To assign cell type annotations, query cells were mapped to an hg38-aligned version of a healthy hematopoiesis reference dataset^57^ using Seurat’s anchor-based reference mapping framework.

Differential accessibility analysis was then performed on *ASXL1*-mutant and wild-type cells across multiple cell-type groupings, including CMP-LMPPs, GMPs, GMPs plus GMP-Neut, and stem and progenitor cells collectively. Differential peaks were annotated with nearest genomic features using ChIPseeker^53^. Initial motif enrichment analysis was performed on differential peaks (adjusted p-value < 0.05) using Signac, with transcription factor motifs obtained from the JASPAR 2020 CORE vertebrate collection^62^. Additional motif enrichment analysis was performed with MEME Suite’s^59^ Simple Enrichment Analysis (SEA) tool^63^ using an EVI1-like ETS family motif derived from MECOM regulatory studies^27^.

## Acknowledgements

The authors thank the following Oregon Health and Science University core facilities for their assistance: OHSU Flow Cytometry and Monoclonal Antibody Shared Resource (RRID:SCR_009974). Short read sequencing assays were performed by the OHSU Massively Parallel Sequencing Shared Resource. We gratefully acknowledge Andrew Adey’s lab at OHSU for providing access to their sequencing equipment, which was used for multiple datasets in this study. This study was funded by a Young Investigator Award from the Edward P. Evans Foundation to TPB, 1R01HL157147-01 from the National Heart, Lung, and Blood Institute to JEM, and funding from the OHSU Knight Cancer Institute Center for Cancer Early Detection and Advanced Research to TPB. The research reported in this publication used computational infrastructure supported by the Office of Research Infrastructure Programs, Office of the Director, of the National Institutes of Health under Award Number S10OD034224. The content is solely the responsibility of the authors and does not necessarily represent the official views of the National Institutes of Health.

Conceptualization, TB, JEM and MH; Methodology, TB, JEM, MH, AQ, MT and HC; Software, SW, TN and TB; Validation, MH, AQ, TB, SW, TN and JM; Formal Analysis, MH, SW, TN, TB, AQ, HC and JEM; Investigation, AQ, MH, ZS, MT, SA and HC; Resources, TB and JEM; Data Curation, SW, TN, MH, AQ, ZS and TB; Writing - Original Draft, TB; Writing - Review & Editing, TB, MH and JEM; Visualization, MH, AQ, SW, TN and TB; Supervision, TB and JEM; Project Administration, TB and JEM; Funding Acquisition, TB and JEM

---

**Supplementary Figure 1.**
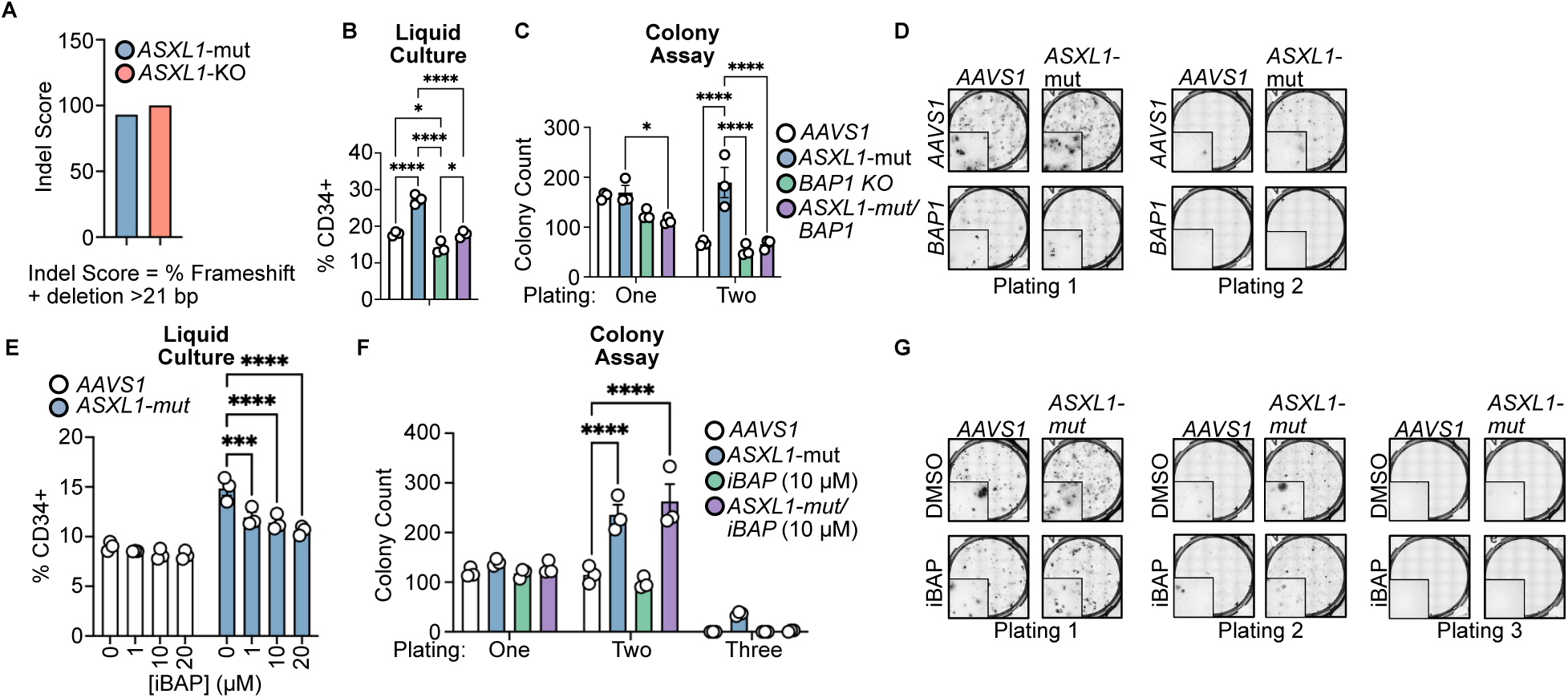
BAP1 is Required for Mutant ASXL1-Driven Oncogenic Phenotypes. (Related to Figure 2) **A.** Results from ICE analysis of CRISPR indels in cord blood CD34+ after Cas9 RNP electroporation and *ASXL1* truncation. Indel score is the % of indels that result in an FS or are >21 bp. **B.** *ASXL1* mutant or *AAVS1*-edited cells with or without BAP1 co-deletion were cultured in multi-lineage differentiation media for 7 days, and CD34 abundance was measured using FACS. **C.** *ASXL1* mutant or *AAVS1*-edited cells with or without BAP1 knockout were plated in methylcellulose and counted by a blinded observer after 14 days. Cells were replated and counted again after 14 days. **D**. Representative Images from **C. E.** *ASXL1* mutant or *AAVS1*-edited cells were treated win increasing concentrations of the BAP1 inhibitor iBAP and cultured in multi-lineage differentiation media for 7 days. CD34 abundance was measured using FACS. **F.** *ASXL1* mutant or *AAVS1*-edited cells were treated with iBAP and plated in methylcellulose and counted by a blinded observer after 14 days. Cells were replated and counted again after 14 days. **D**. Representative Images from **C.** *=p<0.05, ***=p<0.001, ****= p<0.0001 as measured by a two-way ANOVA with a Holm-Sidak post-test.

**Supplementary Figure 2.**
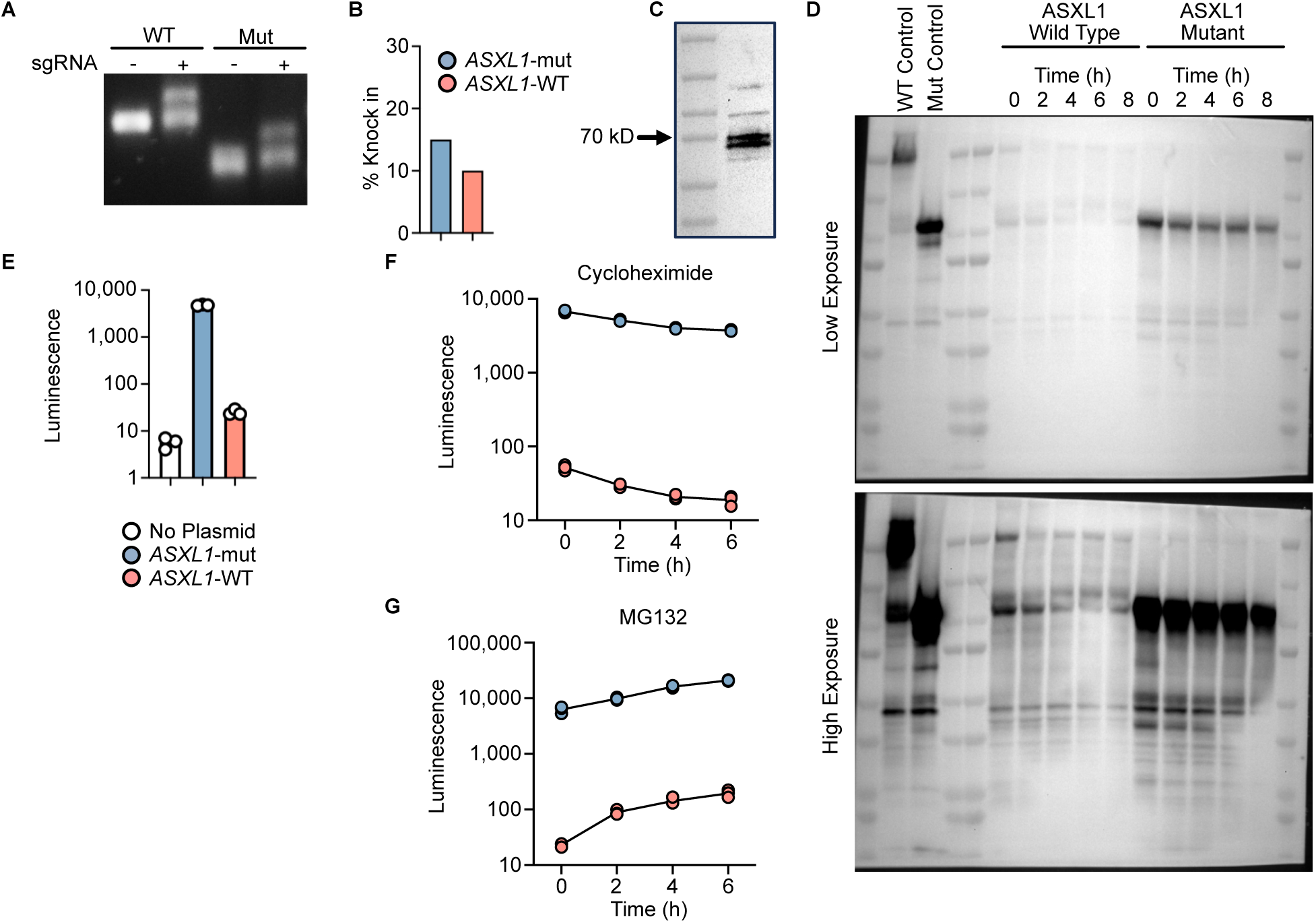
ASXL1 Mutations Drive Protein Stabilization. (Related to Figure 3) **A.** Gel electrophoresis of ICE PCR products. **B.** Quantification of integration efficiency by ICE analysis. **C.** Single-cell cloned U937 cells with the insertion of an ASXL1-truncating HiBIT tag as in Figure 3A. Western blotting confirms protein expression. **D.** Full western blot images from Figure 3C. **E-G.** Un-normalized luminescent signal from Figure 3 D-F.

**Supplementary Figure 3.**
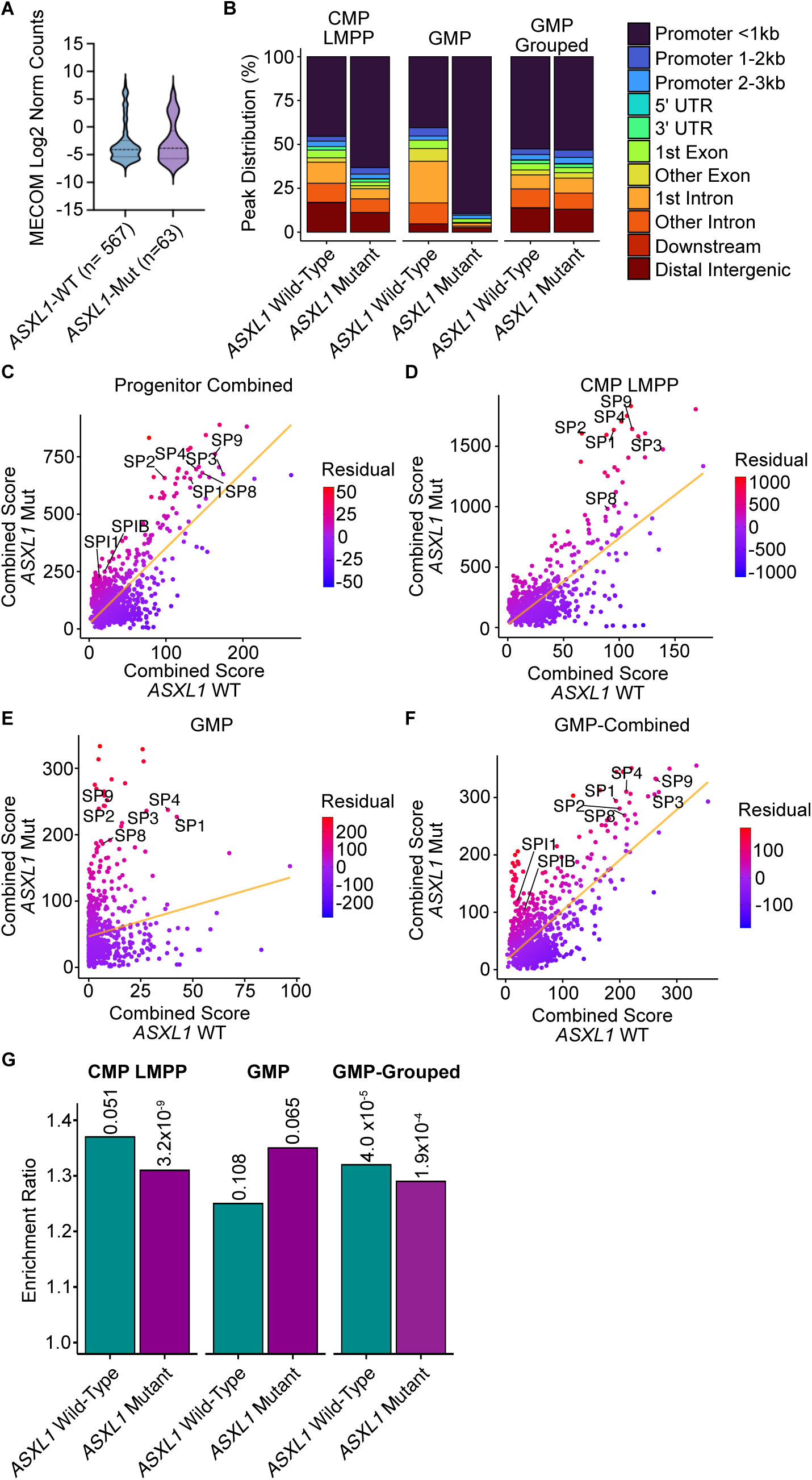
Aberrant MECOM and SP Activity in ASXL1 Mutant AML. (Related to Figure 7) **A.** *MECOM* mRNA expression in *ASXL1*-mutant and wild-type AML. **B.** Peak annotation for differentially accessible regions in *ASXL1* mutant and wild-type cells in the indicated cell types. **C-F.** Database-guided motif enrichment analysis in peaks with increased accessibility in ASXL1 mutant cells (Y-Axis) or ASXL1 wild-type cells (X-Axis). Combined Score = −log10padj x Fold Enrichment for motif enrichment. Points are colored by their distance from the trend line (Residuals). **G.** MECOM motif enrichment within differentially accessible peaks in the indicated cell types. Labels above each bar represent q-values.

